# Evidence for Late Pleistocene origin of *Astyanax mexicanus* cavefish

**DOI:** 10.1101/094748

**Authors:** Julien Fumey, Hélène Hinaux, Céline Noirot, Claude Thermes, Sylvie Rétaux, Didier Casane

**Author notes:** Corresponding author: Didier Casane, Laboratoire Évolution, Génomes, Comportement, Écologie, UMR 9191 CNRS, 1 avenue de la Terrasse, 91198 Gif sur Yvette, France.

## Abstract

**Background:** Cavefish populations belonging to the Mexican tetra species *Astyanax mexicanus* are outstanding models to study the tempo and mode of adaptation to a radical environmental change. They share similar phenotypic changes such as blindness and depigmentation resulting from independent and convergent evolution. As such they allow examining whether their evolution involved the fixation of preexisting standing genetic variations and/or *de novo* mutations. Cavefish populations are currently assigned to two main groups, the so-called "old" and "new" lineages, which would have populated several caves independently and at different times. However, we do not have yet accurate estimations of the time frames of evolution of these populations.

**Results:** First, we reanalyzed the geographic distribution of mitochondrial and nuclear DNA polymorphisms and we found that these data do not support the existence of two cavefish lineages, neither the ancient origin of the “old” lineage. Using IMa2, a program based on a method that does not assume that populations are at mutation/migration/drift equilibrium and thus allows dating population divergence in addition to demographic parameters, we found that microsatellite polymorphism strongly supports a very recent origin of cave populations (*i.e.* less than 20,000 years). Second, we identified a large number of single-nucleotide polymorphisms (SNPs) in transcript sequences of pools of embryos (Pool-seq) belonging to the “old” Pachón cave population and a surface population from Texas. Pachón cave population has accumulated more neutral substitutions than the surface population and we showed that it could be another signature of its recent origin. Based on summary statistics that can be computed with this SNP data set together with simulations of evolution of SNP polymorphisms in two recently isolated populations, we looked for sets of demographic parameters that allow the computation of summary statistics with simulated populations that are similar to the ones with the sampled populations. In most simulations for which we could find a good fit between the summary statistics of observed and simulated data, the best fit occurred when the divergence between simulated populations was less than 30,000 years.

**Conclusions:** Although it is often assumed that some cave populations such as Pachón cavefish have a very ancient origin, within the range of the late Miocene to the middle Pleistocene, a recent origin of these populations is strongly supported by our analyses of two independent sets of nuclear DNA polymorphism using two very different methods of analysis. Moreover, the observation of two divergent haplogroups of mitochondrial and nuclear genes with different geographic distributions support a recent admixture of two divergent surface populations before the isolation of cave populations. If cave populations are indeed only several thousand years old, many phenotypic changes observed in cavefish would thus have mainly involved the fixation of genetic variants present in surface fish populations and within a very short period of time.

## Background

Two well-differentiated morphotypes, surface fish and cavefish, are found in the species *Astyanax mexicanus*. Twenty-nine cavefish populations have been discovered so far in limestone caves in the Sierra de El Abra region of northeastern Mexico [1, 2](**Figure 1**). Cavefish differ from their surface counterparts in numerous morphological, physiological and behavioral traits, the most striking being that most cavefish lack functional eyes and are depigmented [3]. Most caves inhabited by cavefish share a number of abiotic and biotic characteristics such as constant darkness and absence of predators, and most cavefish show evolution of a number of characters [4], either because they are dispensable - regressive traits - such as loss of eyes and pigmentation [5], or because they are involved in the adaptation - constructive traits - to this environment which is inhospitable for most fishes. For example, cavefish have a lower metabolic rate [6–8], produce larger eggs [9], have more and larger superficial neuromasts involved in vibration attraction behavior [10–12], sleep very little [13, 14], have shifted from fighting to foraging behavior [15], have larger numbers of taste buds [16, 17], have enhanced chemosensory capabilities [18] and have enhanced prey capture skill at both the larval and adult stages [11, 19, 20].

**Figure 1.**
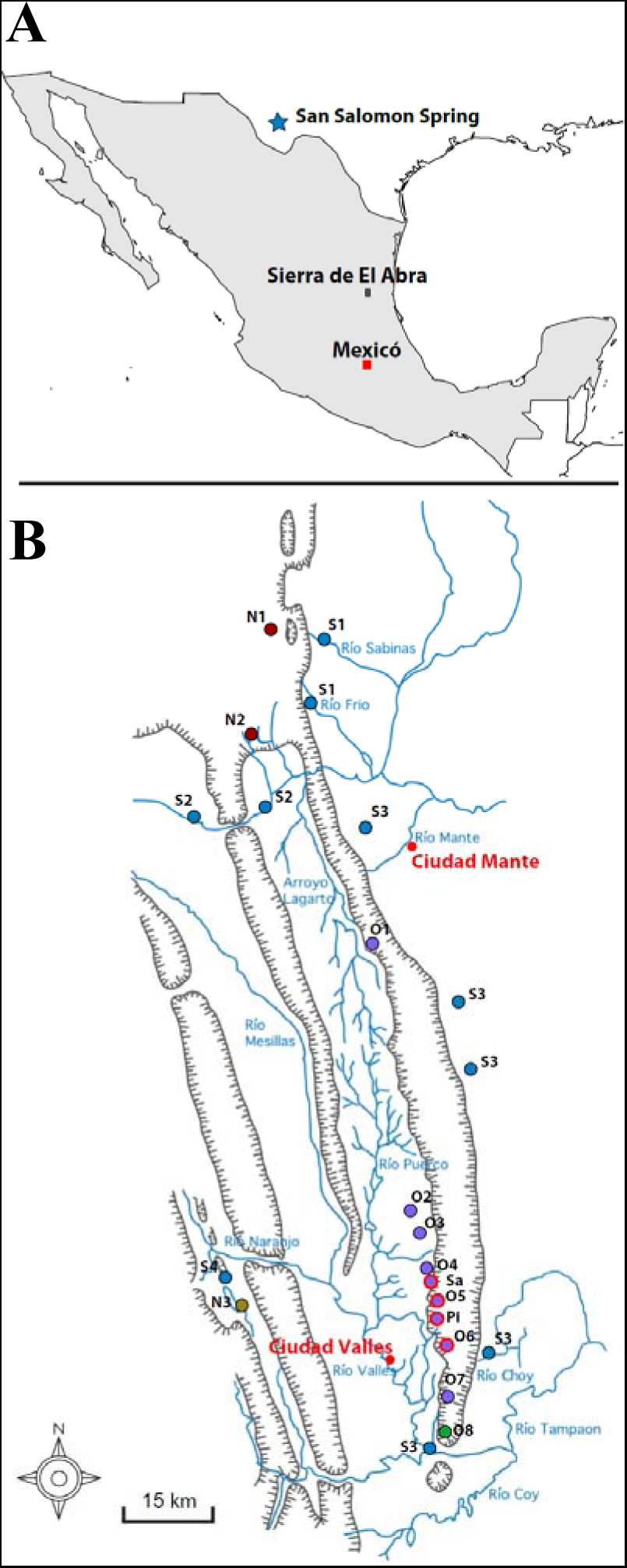
Maps showing cave and surface sampling sites. (A) Sites in Mexico and Texas. (B) Sites in the Sierra de El Abra region in Mexico. Surface fish: S1 to S4. Cave populations: O1 = Pachón, O2 = Yerbaniz, O3 = Japonés, O4 = Arroyo, O5 = Tinaja; O6 = Curva, O7 = Toro, O8 = Chica, N1 = Molino, N2 = Caballo Moro, N3 = Subterráneo (See [33] for a more detailed description of the sampling sites), Sa = Sabinos, Pi = Piedras. Outer circle in red: “G” mtDNA.

Very significant advances have been made in identifying proximal mechanisms [21], that is mutations that have changed physiological, developmental, and behavior traits of cavefish and new molecular tools available today will allow us to identify such mutations at an ever increasing pace [22–26]. However it is much more tricky to disentangle distal mechanisms [21], *i.e.* evolutionary mechanisms. Were these mutations already present at low frequency in surface fish standing variation or did they appear after settlements? Are pleiotropic effects and epistatic interactions important in these evolutionary processes? What is the impact of recombination, genetic drift, selection and migration in cavefish evolution? These questions have fueled discussions on the relative importance of these different evolutionary mechanisms [12, 17, 27–31].

In order to analyze several of these issues such as the relative weight of selection, migration and genetic drift, it would be very useful to have accurate estimations of some demographic and population genetic parameters to describe the dynamic of cavefish evolution. Gene flow from the surface populations has been estimated to be from very low, if any, to very high, depending on the cave population examined. Some studies have also found significant and higher gene flow from cave to surface populations than in the opposite direction [32–37]. Moreover, as some caves are very close to each other, fish migrations within cave clusters are likely.

Among model parameters particularly important to describe the evolution of a cavefish population are: 1) the time at which settlement occurred and 2) how long it took for surface fish to adapt to the cave environment. As shown below, no reliable ages were available but *Astyanax mexicanus* cave populations have nevertheless been assigned to two groups, the so-called “old” and “new” lineages, which would have populated several caves independently and at different times [37–39], reviewed in [2]. However, the age of cavefish settlement has been estimated for two populations only, those inhabiting the Pachón and Los Sabinos caves, which both belong to the “old” lineage. On the basis of allozyme polymorphism [32] and a population genetic method specifically designed to estimate the time after divergence between incompletely isolated populations of unequal sizes (such as cave and surface populations), these populations were estimated to be 710,000 and 525,000 years old, respectively, suggesting that they could be ancient [40]. However, the small number of loci studied at that time (17 allozyme loci scored), the absence of polymorphism in Pachón and very low polymorphism in Los Sabinos did not allow accurate estimations. The standard error (SE) was very large, 460,000 and 330,000 years, respectively. Assuming a normal distribution [41], the 95% confidence interval is ± 1.96 × SE. It implies that these populations could be either very recent (a couple of thousand years and even less) or very ancient (about 1,5 million years). Based on this analysis, the only safe conclusion is that these cave populations are not millions of years old. The large uncertainty associated to these estimations is probably the reason why they are rarely cited by investigators working on these cavefish.

The hypothesis of a very ancient origin of the “old” cavefish lineage, *i.e.* millions years ago, relies only on discussions of mitochondrial DNA (mtDNA) phylogenies of surface fish and cavefish, showing two highly divergent mitochondrial haplogroups [37, 39, 42]. However, ancient coalescence of mtDNA haplogroups does not necessarily imply an ancient isolation of some cave populations, *i.e.* the time of separation of the populations is not necessarily equal, not even close, to the time of coalescence of the mtDNA sequences. An alternative hypothesis that would lead to the same observation is a recent admixture of two divergent surface populations followed by one or several fish settlements in caves. These hypotheses will be tested below.

Nevertheless, assuming an ancient origin of cave populations and thus that surface and cave populations are at mutation/migration/drift equilibrium, estimation of differentiation [32–34, 38] and migration rates among populations [33, 35] were performed using microsatellite polymorphism. More recently a phylogenetic analysis was performed using a large SNP data set in order to estimate the number of independent cave settlements, but this approach did not allow dating as constant sites were discarded [43]. Moreover, this analysis did not support the two lineage hypothesis.

In summary, and except an attempt using an allozymes data set unfortunately too small to give accurate estimations, no dating has ever been performed using nuclear markers to directly test the assumption that some cavefish populations are millions years old.

Several observations led us to doubt about its accuracy. In particular, looking at sequences of Pachón cavefish, we did not find many obvious loss-of-function mutations, such as frameshifts and stop codons, in eye-specific crystallin genes [26] and opsin genes [44–46](unpublished results), an unexpected observation if this population was established for several hundred thousand years, and which becomes very unlikely if it was established more than one million years ago [47]. Indeed, other fish that could be confined into caves for millions of years have fixed loss-of-function mutations in several opsins and crystallins genes [48–50].

Here, we analyzed published sequence data sets and we found that different nuclear loci have different phylogenies that are not congruent with the mtDNA phylogeny. Moreover, using a published microsatellite data set and an approach that allow dating the isolation of closely related populations when there is gene flow, we obtained good evidence of a recent origin of all cavefish population analyzed, *e.g* likely less than 20,000 years ago, notwithstanding their “old” or “young” classification. In these analyses, estimations of effective population sizes and migration rates were more coherent with expectations than in previous analyses: effective population sizes for cavefish were at least one order of magnitude smaller than those for surface fish; gene flows were from the surface to caves and not the other way around. In order to corroborate these novel estimations with an independent data set and using a very different method, we identified and analyzed a large number of single-nucleotide polymorphisms (SNPs) in transcript sequences of two pools of embryos (Pool-seq) belonging to the Pachón cave population and a surface population from Texas. The comparison of these data with simulations suggests that the Pachón cave population has probably been underground less than 30,000 years.

Both dating methods gave congruent estimations of the age of the Pachón cave population, and they pointed to a very recent origin.

The new time frame we propose for the evolution of *A. mexicanus* cavefish suggests that the many phenotypic changes observed in these cavefish may have mainly involved the fixation of genetic variants present in surface fish populations, and within a very short period of time.

## Results

### Nuclear haplotype phylogenies

There are currently few nuclear genes for which sequences from different cave and surface fish have been published. A fragment of the coding sequence of Rag1 is available for a large sample of surface *Astyanax* spp. but only one cavefish [42]. A phylogeny based on this gene alone has not been published. We performed this phylogenetic analysis **(Additional File 1; Figure S1)**. This phylogeny is so very poorly resolved that it cannot support the mtDNA phylogeny **(Additional File 1; Figure S2)** nor any another phylogeny. Then we analyzed a series of genes (Mc1r, Oca2, Mc4r, Mc3r, Lepb, Lepr, Pomcb) for which at least four sequences were available, three from cavefish and one from a surface fish **(Additional File 2)**. For Mc4r, Mc3r, Lepb, Lepr, Pomcb, sequences were obtained for the same localities (Surface, Pachón, Tinaja and Molino) [51] **(Figure 1)**. The matrix of informative sites in each gene is shown (**Figure 2A**). When more than one informative site was found within a gene, all informative sites supported the same tree topology but different genes supported different tree topologies. Overall the three possible unrooted trees were similarly supported **(Figure 2B, Additional File 2)**.

**Figure 2.**
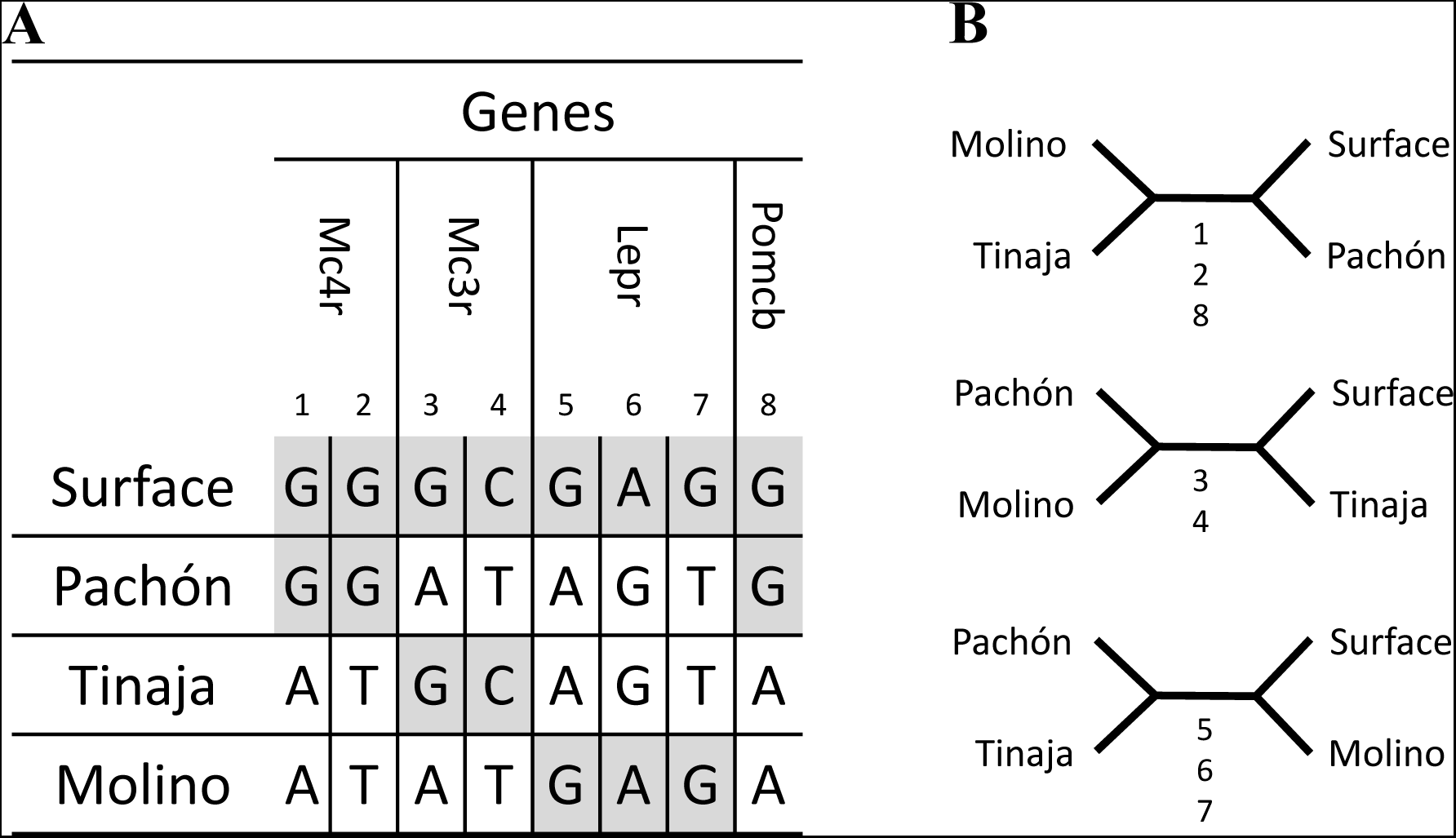
Incongruence of four taxon nuclear gene phylogenies. (A) Informative sites in Mc4R, Mc3R, LeptinR and PomC_B. (B) Unrooted trees with four taxa. Informative sites equally support the three possible trees.

Partial or complete coding sequences of five other genes (Per1, Per2, Tef1, Cry1a and Cpd photolyase) from three localities (Pachón and Chica caves and a surface locality close to Micos) were also available [52]. These sequences were aligned with the sequence of the Texas surface fish and the most parsimonious unrooted tree was reconstructed for each gene. Despite the fact that two divergent haplotypes were found for some genes, the two surface fish sequences were always very close **(Additional File 2)**. In sum, the incongruence of the phylogenies of these genes suggests that they have independent evolutionary history.

### Dating with microsatellites

We next re-analyzed a previously published data set of microsatellite polymorphisms [33] using IMa2 [53]. This program, which implements a method based on simulations of coalescence of samples of alleles, allows the estimation of the marginal posterior probability density of population sizes, migration rates and divergence times. We performed a series of pairwise analyses involving a cave population and a surface population. We also analyzed the divergence of three cave populations. The estimated marginal posterior probability densities of the model parameters obtained with Pachón (O1) and a surface population (S3), using 22 loci and a random sample of 60 alleles per locus per population are shown (**Figure 3**). Similar distributions were obtained using the first half and the second half of the MCMC chain, suggesting that the sampling process has been long enough to get stable posterior probability distributions. Moreover, these posterior distributions have a single sharp peak and the probabilities are low for the extremes values of the prior distributions, suggesting that the selection of the maximum value of each parameter was suitable. In IMa2, demographic parameters are scaled to the mutation rate. Estimations of these parameters thus depends on prior on the mutation rate. Assuming that the mutation rate of the microsatellites is 5 × 10^−4^ [33, 54, 55], estimations of population sizes, migration rates and divergence time can be obtained (**Table 1**). The effective population size (Ne) was 150 [150 – 1150] and 10,750 [6,650 – 16,350] for Pachón cave population and surface fish population respectively (the maximum likelihood value is given with the 95% highest posterior density interval between brackets). The ancestral effective population size was 44,850 [32,350 – 73,250]. The divergence time was 5,110 years [1,302 – 18,214]. The migration rates were very low, but about 100 times higher from the surface than from the cave.

**Figure 3.**
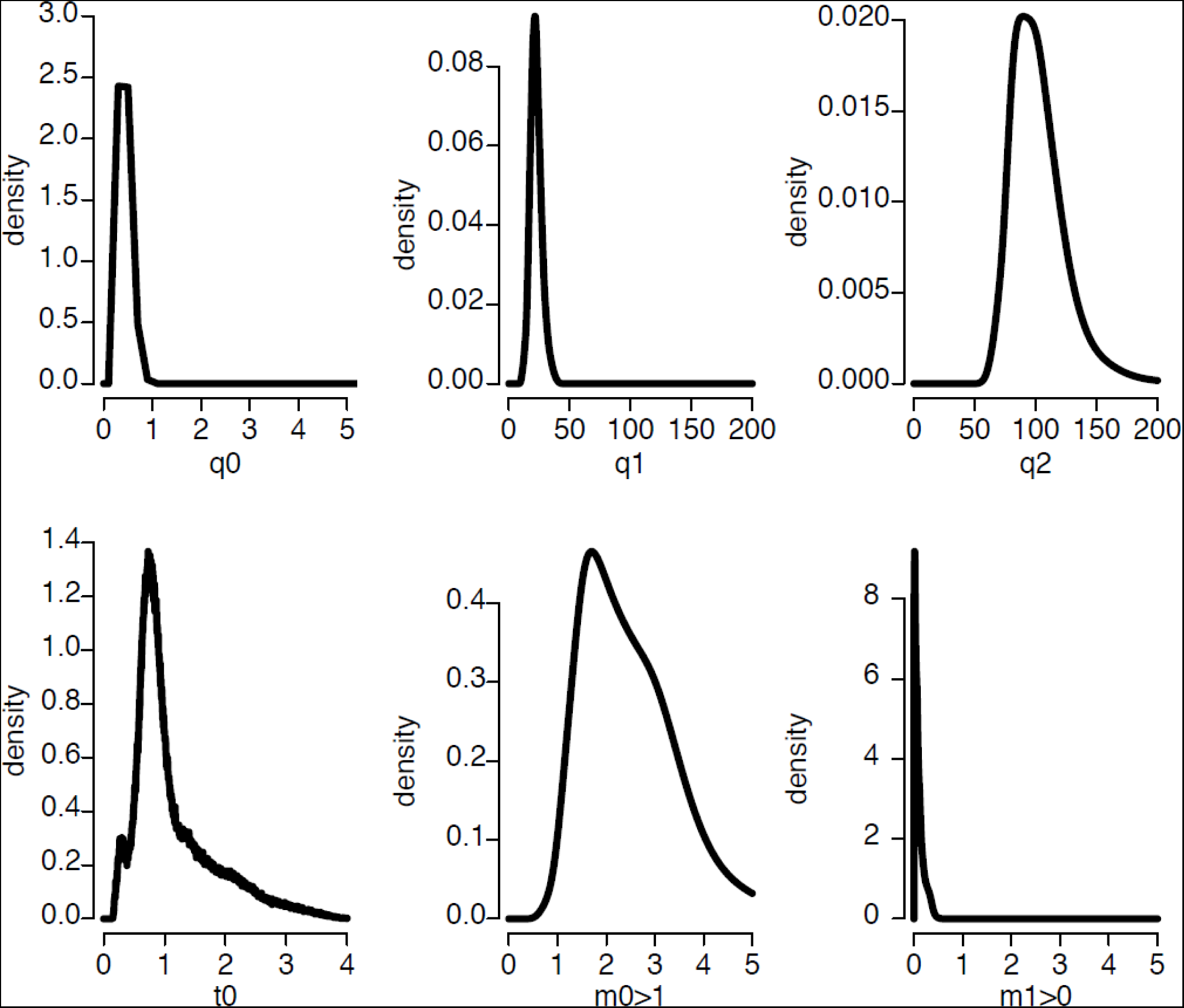
Posterior marginal density plots of the demographic parameters estimated from the isolation with migration model of Pachón cavefish and N3_surface fish. q0 = 4N_0_μ (where N_0_ is the effective size of population 0 [*i.e.* Pachón cavefish] and μ is the mutation rate), q1 = 4N_1_μ (where N_1_ is the effective size of population 1 [*i.e.* N3_Surface fish]), q2 = 4N_2_μ (where N_2_ is the effective size of population 2 [*i.e.* ancestral population]), t0 = tμ/g (where t is the number of generation since divergence and g is the generation time), m0>1 (μ-scaled migration rate from population 0 to population 1 backward in time = μ-scaled migration rate from population 1 to population 0 forward in time), m1>0 (μ-scaled migration rate from population 1 to population 0 backward in time = μ-scaled migration rate from population 0 to population 1 forward in time). Command line: ./IMa2 –iinfileO8S3_n60.u - ooutput_O1S3_n60_longrun.out -b200000 -l100000 -d100 –m10 -q200 -t4 -c1 -p23567 -r25 - u1 -hfg -hn50 -ha0.999 -hb0.3

**Table 1.**
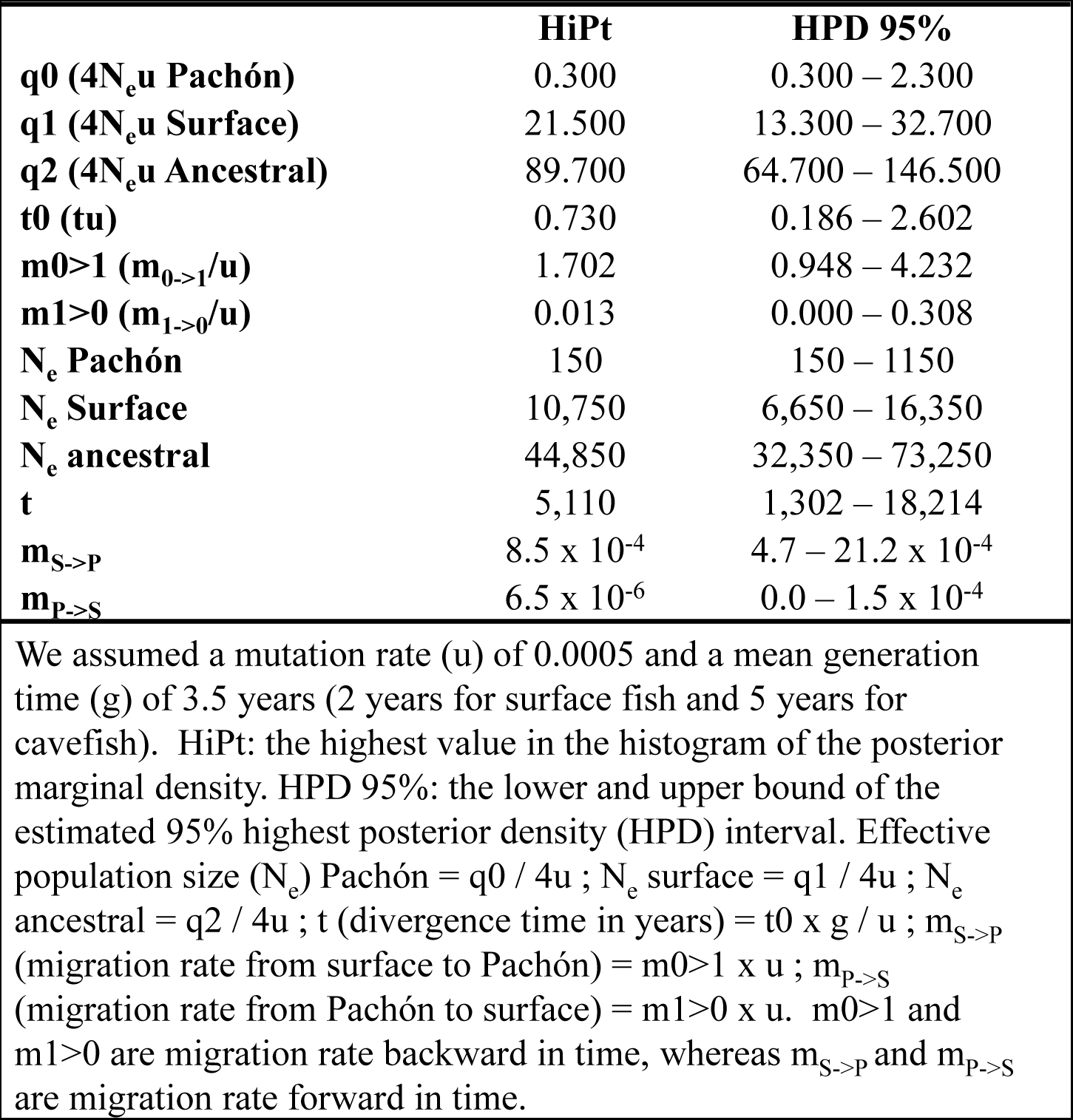
Estimated of demographic parameters with IMa2

|  | **HiPt** | **HPD 95%** |
| --- | --- | --- |
| **q0 (4N<sub>e</sub>u Pachón)** | 0.300 | 0.300 – 2.300 |
| **q1 (4N<sub>e</sub>u Surface)** | 21.500 | 13.300 – 32.700 |
| **q2 (4N<sub>e</sub>u Ancestral)** | 89.700 | 64.700 – 146.500 |
| **t0 (tu)** | 0.730 | 0.186 – 2.602 |
| **m0>1 (m<sub>0->1</sub>/u)** | 1.702 | 0.948 – 4.232 |
| **m1>0 (m<sub>1->0</sub>/u)** | 0.013 | 0.000 – 0.308 |
| **N<sub>e</sub> Pachón** | 150 | 150 – 1150 |
| **N<sub>e</sub> Surface** | 10,750 | 6,650 – 16,350 |
| **N<sub>e</sub> ancestral** | 44,850 | 32,350 – 73,250 |
| **t** | 5,110 | 1,302 – 18,214 |
| **m<sub>S->P</sub>** | 8.5 x 10<sup>-4</sup> | 4.7 – 21.2 x 10<sup>-4</sup> |
| **m<sub>P->S</sub>** | 6.5 x 10<sup>-6</sup> | 0.0 – 1.5 x 10<sup>-4</sup> |

We assumed a mutation rate (u) of 0.0005 and a mean generation time (g) of 3.5 years (2 years for surface fish and 5 years for cavefish). HiPt: the highest value in the histogram of the posterior marginal density. HPD 95%: the lower and upper bound of the estimated 95% highest posterior density (HPD) interval. Effective population size (N<sub>e</sub>) Pachón = q0 / 4u ; N<sub>e</sub> surface = q1 / 4u ; N<sub>e</sub> ancestral = q2 / 4u ; t (divergence time in years) = t0 x g / u ; m<sub>S->P</sub> (migration rate from surface to Pachón) = m0>1 x u ; m<sub>P->S</sub> (migration rate from Pachón to surface) = m1>0 x u. m0>1 and m1>0 are migration rate backward in time, whereas m<sub>S->P</sub> and m<sub>P->S</sub> are migration rate forward in time.

With the same sample size (*i.e.* number of loci and number of alleles / locus / population), the same analyses were performed with Chica cave (O8) and the same surface population (S3) **(Additional File 1; Figure S3)** and with Pachón (O1) and Chica (O8) caves **(Additional File 1; Figure S4)**. Such a large number of alleles for a large number of loci was not available for other useful analyses **(Additional File 1; Table S1)**, we thus used smaller samples. With a sample of 14 loci and 40 alleles per locus per population, the same analysis was performed with Molino cave (N1) and a surface population (S1) **(Additional File 1; Figure S5)**. With a sample of 18 loci and 40 alleles per locus per population, the same analysis was performed with Caballo Moro cave (N2) and a surface population (S2) **(Additional File 1; Figure S6)**. With a sample of 21 loci and 40 alleles per locus per population, the same analysis was performed with Subterráneo cave (N3) and a surface locality (S4) **(Additional File 1; Figure S7)**. With a sample of 14 loci and 20 alleles per locus per population, the same analysis was performed with Curva cave (O6) and a surface locality (S3) **(Additional File 1; Figure S8)**, and with Curva cave (O6) and Pachón cave (O1) **(Additional File 1; Figure S9)**. The results of these analyses are highly consistent (**Table 2**). Cave effective population size (Ne) was always low, a few hundreds. Extant surface population Ne was consistently found to be about 10,000 whatever the sampling localities. The ancestral surface population Ne was consistently found larger than the extant surface population Ne, *i.e.* between 2 and 4 times larger. The migration rates were always low. The maximum likelihood of the divergence time was similar in all cases, in the range of 1,000 to 10,000 years, the posterior distributions largely overlapping.

**Table 2.** Estimated of demographic parameters with IMa2

| Populations |  | **p0 – p1**<br>**O1 –S3** | **p0 – p1**<br>**O6 –S3** | **p0 – p1**<br>**O8 –S3** | **p0 – p1**<br>**O1 –O6** | **p0 – p1**<br>**O1 –O8** | **p0 – p1**<br>**N1 –S1** | **p0 – p1**<br>**N2 –S2** | **p0 – p1**<br>**N3 –S4** |
| --- | --- | --- | --- | --- | --- | --- | --- | --- | --- |
| **N<sub>e</sub> p0** | HiPt | 150 | 50 | 250 | 250 | 250 | 150 | 850 | 450 |
|  | HPD 95% | [150 – 1150] |  | [150 – 1,250] | [150 – 1,250] | [150 – 1150] | [0 – 950] | [350 – 1,850] | [250 – 1,650] |
| **N<sub>e</sub> p1** | HiPt | 10,750 | 13,550 | 5,850 | 350 | 550 | 10,550 | 3,850 | 6,950 |
|  | HPD 95% | [6,650 – 16,350] |  | [1,950 – 60,350] | [150 – 1,350] | [150 – 1,550] | [6,050 – 15,950] | [2,150 – 6,950] | [1,550 – 14,350] |
| **N<sub>e</sub> ancestral** | HiPt | 44,850 | 33,850 | 42,450 | 24,050 | 26,650 | 47,050 | 37,950 | 43,350 |
|  | HPD 95% | [32,350 – 73,250] |  | [28,750 – 62,650] | [11,950 – 50,750] | [14,150 – 44,150] | [28,950 – 81,550] | [25,250 – 65,350] | [20,350 – 69,150] |
| **t** | HiPt | 5,110 | 2,982 | 798 | 2,540 | 1,274 | 10,150 | 4,074 | 770 |
|  | HPD 95% | [1,302 – 18,214] |  | [30 – 8,302] | [860 – 5,980] | [266 – 2,422] | [2,002 – 24,990] | [1,834 – 10,626] | [406 – 17,234] |
| **m<sub>1->0</sub>** | HiPt | 8.5 x 10<sup>-4</sup> | 27.9 x 10<sup>-4</sup> | 11.2 x 10<sup>-4</sup> | 8.6 x 10<sup>-4</sup> | 3.4 x 10<sup>-4</sup> | 4.9 x 10<sup>-4</sup> | 3.0 x 10<sup>-4</sup> | 29.0 x 10<sup>-4</sup> |
|  | HPD 95% | [4.4 – 21.2 x 10<sup>-4</sup>] |  | [6.4 – 50.0 x 10<sup>-4</sup>] | [0.5 – 24.5 x 10<sup>-4</sup>] | [0.0 – 8.7 x 10<sup>-4</sup>] | [1.0 – 15.2 x 10<sup>-4</sup>] | [0.0 – 8.3 x 10<sup>-4</sup>] | [9.0 – 57.2 x 10<sup>-4</sup>] |
| **m<sub>0->1</sub>** | HiPt | 6.5 x 10<sup>-6</sup> | 4.6 x 10<sup>-4</sup> | 7.93 x 10<sup>-4</sup> | 2.5 x 10<sup>-6</sup> | 4.3 x 10<sup>-4</sup> | 5.0 x 10<sup>-6</sup> | 1.5 x 10<sup>-6</sup> | 10.2 x 10<sup>-4</sup> |
|  | HPD 95% | [0.0 – 1.5 x 10<sup>-4</sup>] |  | [1.3 – 10.7 x 10<sup>-4</sup>] | [0.0 – 8.5 x 10<sup>-4</sup>] | [0.0 – 10.1 x 10<sup>-4</sup>] | [0.0 – 1.3 x 10<sup>-4</sup>] | [0.0 – 1.8 x 10<sup>-4</sup>] | [3.8 – 17.2 x 10<sup>-4</sup>] |

See Table 1 for definition of parameters and Figure 1 for localization of populations

These results suggest that all the cave populations analyzed are very recent, regardless of their classification as ancient, recent, isolated or mixed, and regardless of their mtDNA haplotype.

### SNPs and substitution rates in surface and cave populations

Next, we set out to corroborate these previous findings with an independent method, through the analyses of SNPs found in surface fish (SF) and Pachón cavefish (CF) transcriptomes, coupled to simulations. We used transcriptome sequence data sets from pooled embryos (**Additional File 1; Figure S10**). We defined eight classes of polymorphic sites according to the presence of an ancestral and/or a derived allele in SF and CF populations, using the Buenos Aires tetra (*Hyphessobrycon anisitsi*) as an outgroup (**Figure 4**).

**Figure 4.**
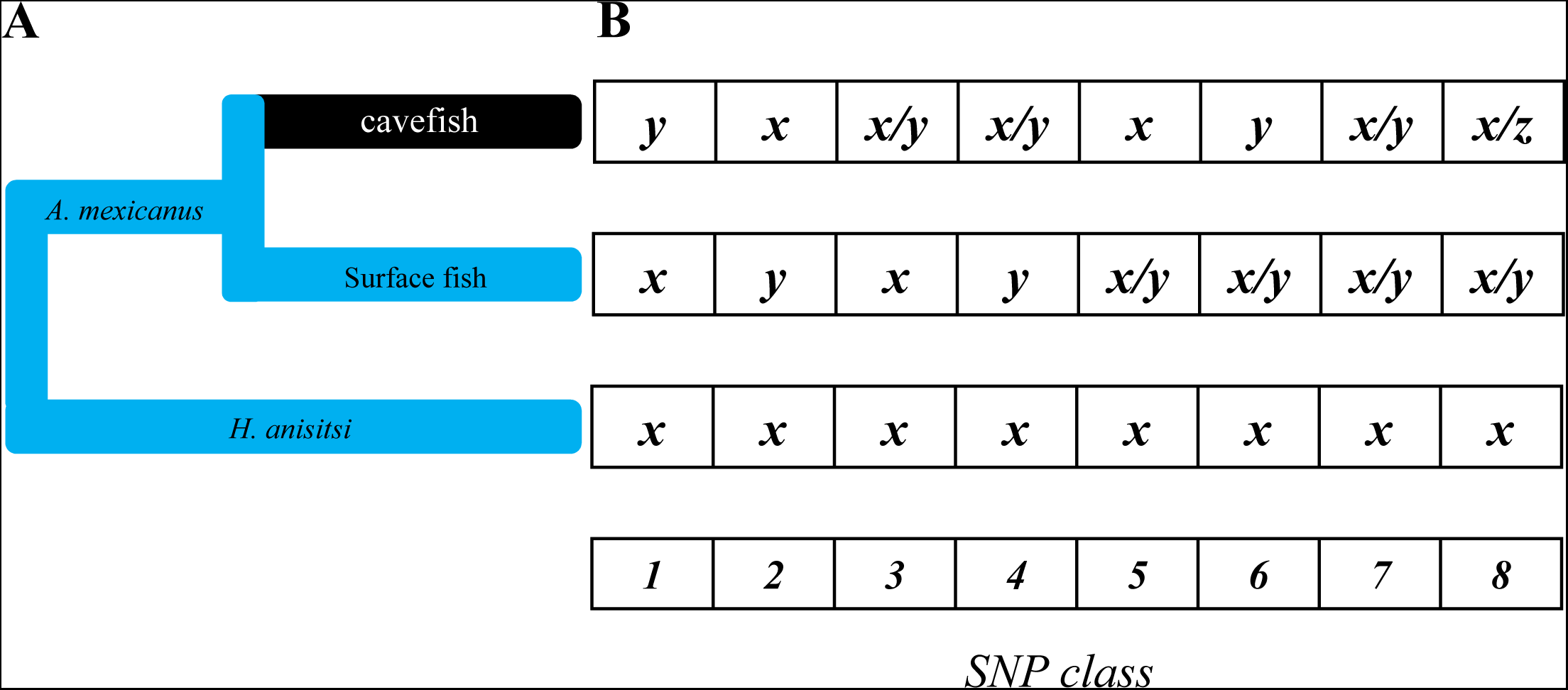
Analysis of polymorphism in *Astyanax mexicanus* Texas surface *vs* Pachón cave population, using *Hyphessobrycon anisitsi* as outgroup. (A) Evolutionary model. (B) The eight SNP classes correspond to the polymorphism patterns that can be found within and between two populations. Class 1: Different fixed alleles in each population, derived allele in cavefish; Class 2: Different fixed alleles in each population, derived allele in surface fish; Class 3: Polymorphism in cavefish, ancestral fixed allele in surface fish; Class 4: Polymorphism in cavefish, derived fixed allele in surface fish; Class 5: Polymorphism in surface fish, ancestral fixed allele in cavefish; Class 6: Polymorphism in surface fish, derived fixed allele in cavefish; Class 7: Shared polymorphism; Class 8: Divergent polymorphism. x, y and z can be one of the four nucleotides A, T, G, C.

We estimated the frequencies of these eight SNP classes at synonymous, non-coding and non-synonymous sites (**Table 3**). The frequencies of SNPs in the eight classes were robust according to different parameter thresholds used to include SNPs in the analysis (**Materials and methods**, **Additional File 1; Figure S11, Table S2a and Table S2b)**. The ratio (SF/CF) of synonymous, non-coding and non-synonymous polymorphism was 3.08, 2.71 and 2.34, respectively, and the ratio (CF/SF) of derived fixed alleles was 2.34, 1.45 and 1.52, respectively. These results indicated that the level of polymorphism was higher in the SF population, but the number of fixed derived alleles was higher in the CF population.

**Table 3.**
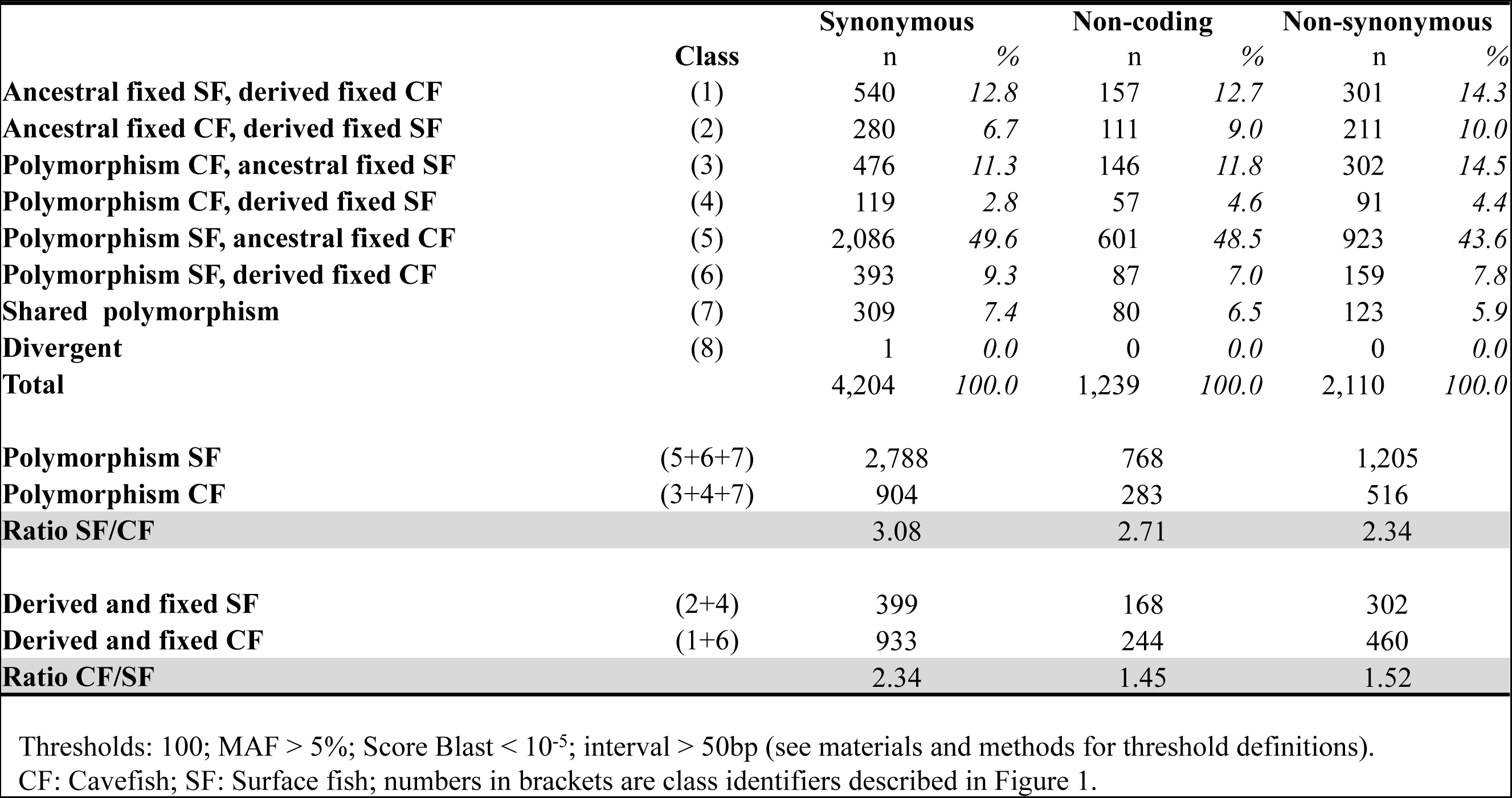
Classification of polymorphisms in *Astyanax mexicanus* Texas surface *vs* Pachón cave populations

### Dating with SNPs

In order to make an estimation of the age of the Pachón cave population with the SNPs that is independent of the estimation made with microsatellite polymorphism, we compared the observed summary statistics of synonymous polymorphism with the summary statistics of neutral polymorphism in simulated populations using the model of evolution implemented in IMa2, *i.e.* an ancestral population divided into two populations at some point in the past. Each population had its own effective population size (**Figure 4**). Migrations between populations were allowed. Whereas IMa2 is based on simulation of coalescence of a sample of alleles (backward in time), we did simulations forward in time of genetic drift of allele frequencies in whole populations, which was more adapted to analyze our SNP dataset. Both approaches have their own advantages and limits [56]. Simulations of genetic drift in two populations recently and partially isolated allowed to estimate summary statistics that could be easily compared with the observed summary statistics. In addition, the implementation of our own simulation program allowed to take into account a change of generation time in cavefish and to compute more easily the evolution of summary statistics through time.

We tested several wide ranges of parameters values (population sizes, migration rates, isolation time) that were defined taking into account the results obtained with IMa2, previous populations genetics studies and information gathered through several trips to the Pachón cave.

A few set of parameters allowed a good fit between the summary statistics of the observed and simulated polymorphism (**Additional File 1; Table S3a, Table S3b, Table S3c and Table S3d**). As an example, we can consider a case, *i.e.* a parameter set, which gave a good fit (in **Additional File 1; Table S3a**, framed in yellow). In this simulation the ancestral population size was set to 10,000 and was at mutation/drift equilibrium; after the separation of the surface and cave populations, the cave population size was set to 1,250 and the Texas surface population size was set to 10,000; the probability of migration per year from surface to cave was 0.001 and the number of migrants was 1% of the cave population size (*i.e.* 12 fish); the generation time of the cavefish was set to 5 years and the generation time of the surface fish was set to 2 years. Every 100 years (i.e. 50 SF generations, or 20 CF generations), 10 fish were sampled in each population to simulate the sampling process when the lab populations were established. Each lab population was then set with a constant effective population size of 10 over 10 generations. Then we compared the frequency of each SNP class in the simulated lab populations with the observed frequency. It was thus possible to check the fit of the summary statistics through time using a goodness of fit score. In this simulation, the best fit (the lowest value of the score) occurred when the age of the cave population was 21,500 years (**Figure 5A**). All SNP class frequencies in the simulated populations fit well (goodness of fit score = 0.68) with the observed frequencies (**Figure 5B**). Then, the older was the divergence of the populations and the worse was the fit (**Figure 5A**). In this simulation, as well as in all other simulations, the mutation rate per generation (u), that is the probability of appearance of a new allele at a new locus in one haploid genome at a given generation, was set to 2.10^−2^. The number of new SNPs that appeared per generation in a population of size N was 2Nu, each with a frequency of 1/2N. This means that in the surface population there was 2 × 10,000 × 2 × 10^−2^ = 400 new SNPs at each generation, and that these 400 new SNPs appeared with an initial frequency of 1 / (2 × 10,000) = 5 × 10^−5^. In parallel, 50 new SNPs appeared with initial frequency of 4 × 10^−4^ in the cave population at each generation. All loci were independent. It is noteworthy that the fit of the actual and simulated polymorphism did not depend on the mutation rate because we compared the relative frequencies of SNP classes rather than their absolute numbers. In other words, if the mutation rate was higher, the number of SNPs in each class was higher, but the relative frequency of each class remained the same. Thus the score of goodness of fit did not depend on the mutation rate. The mutation rate we used was a trade-off between the accuracy of the SNP class frequency estimations in the simulated populations and the time to run a simulation (the higher the mutation rate, the higher the number of polymorphic sites for which allele frequency evolution was simulated). Finally, the estimation of the age of the cave population depends on the generation time in each population as this age, in years, is the product of the number of generation multiplied by the generation time.

**Figure 5.**
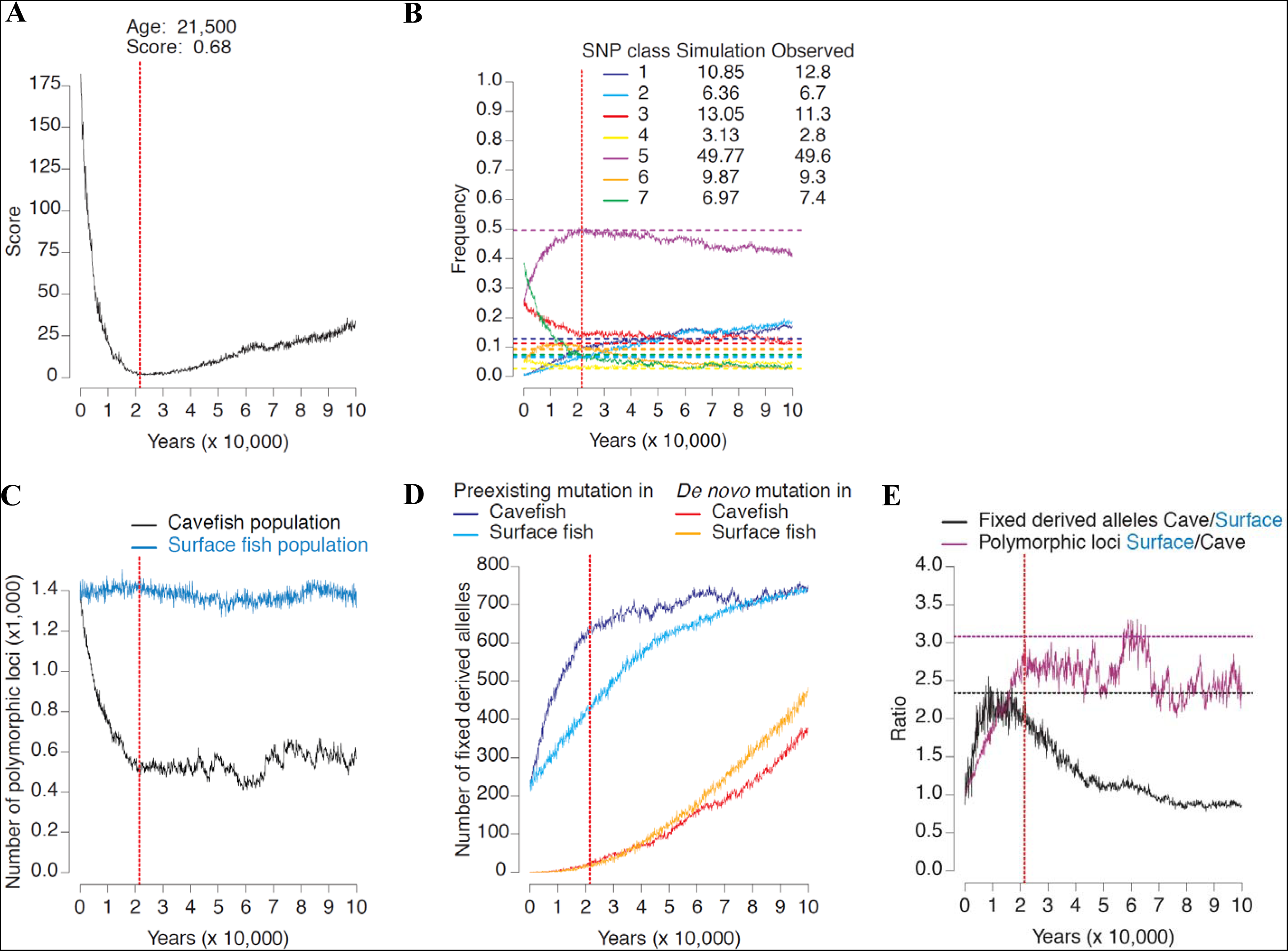
Goodness of fit to the data. The model parameters are: SF population size = 10,000; CF population size = 625; % migrants from surface to cave = 0.1; migration rate from surface to cave = 0.001 / year; SF generation time = 2 years; CF generation time = 5 years; lab population parameters: 10 fish, 10 generations. All the other parameters were set to zero. (A) Score of goodness of fit according to the age of the cave population (t3), the best fit is when the cavefish population is 25,500 years old. (B) Evolution of the SNP class frequencies during the simulation. Horizontal dotted lines are the observed SNP class frequencies. Observed and simulated frequencies at the age of the best fit are shown in the top right corner. (C) Evolution of the number of polymorphic sites in SF and CF during the simulation. (D) Evolution of the number of derived alleles that were fixed in SF and CF during the simulation. (E) Evolution of the SF/CF polymorphism ratio and the CF/SF derived allele ratio that reached fixation during the simulation. Horizontal dotted lines are the observed ratios. The vertical dotted line is the age of the cavefish population for which the best fit was observed.

As the analyses with IMa2 suggested that the ancestral population size was larger than the extant surface population size, we tested ancestral population in the range of 10,000 to 100,000. Surface population size was set in the range of 5,000 to 20,000. Cavefish population size was set in the range of 75 to 10,000. We also took into account migration from the surface to the cave: the probability of migration varied between 0.1 and 0.0001 per year and the percentage of surface fish that migrated into the cave varied between 1% and 0.01% of the cavefish population size. We considered that the migration rate and the number of migrants at each migration from the cave to surface was negligible as it is suggested by the analysis with IMa2 and our observations in the field.

In summary, good fits could be found between observed and simulated summary statistics when the effective population size of the cave population was much smaller than the surface effective population size and when the divergence was recent, in most cases around 20,000 years (**Additional File 1; Table S3a, Table S3b, Table S3c and Table S3d**).

## Discussion

### Evidence against the existence of old and new lineages of *A. mexicanus*

We discuss below that the presence of two divergent mtDNA haplotypes is not *per se* a strong support for the existence of two fish lineages. Moreover, the incongruence of the mtDNA phylogeny with phylogenies obtained with several independent nuclear loci definitively invalidates this hypothesis.

A widely accepted scenario in the community working on *A. mexicanus* cavefish is that some cave populations are ancient, *i.e.* hundreds of thousands or even millions of years old, and related to extinct surface fish, whereas other cave populations are more recent and related to extant surface fish. We demonstrate below that this hypothesis relies only on the existence of two divergent mtDNA haplogroups that are supposed to reflect the existence of two divergent fish lineages. The hypothesis that cavefish originated from two separate surface fish stocks was first formulated on the basis of a NADH dehydrogenase 2 (ND2) phylogeny of cave and surface fish [39]. On the one hand all surface fish from the Sierra de El Abra belonged to a haplogroup named “lineage A”, as well as two surface fish from Texas and a surface fish from the Coahuila state, in northeastern México. Pachón and Chica cavefish also belonged to this haplogroup A. On the other hand, Curva, Tinaja and Sabinos cavefish, living in caves that are geographically close to each other, belonged to another and well differentiated haplogroup named “lineage B”. The authors concluded that the cavefish belong to an old stock of fish, “the lineage B” that was present at the surface a long time ago, but now extinct and replaced by surface fish with haplotypes belonging to haplogroup A. Noteworthy this hypothesis implies that the mtDNA haplotype A1 found in Pachón and Chica cavefish (a haplotype found in most surface localities) is the result of recent mtDNA introgressions into these caves. The authors of this publication proposed another explanation: whereas it is likely that introgression can occur in Chica cave where surface fish and hybrids have been found, they suggested that Pachón cavefish, that seem much more isolated, have evolved independently and more recently than haplogroup B cavefish, and they are undergoing troglomorphic evolution more rapidly than other cavefish populations [39]. These hypotheses were among the most parsimonious that could be formulated at that time with this data set. It is important to note that, and even if we accept the *ad hoc* hypothesis that a surface population has been replaced by another surface population with mtDNAs belonging to a divergent haplogroup, it does not necessarily imply that the age of the cavefish related to the first population is the time of coalescence of the two divergent mtDNA haplogroups. Indeed, two surface populations could have evolved independently for a very long time in two separated Mexican regions, allowing the evolution of two divergent haplogroups, but one population (the extant population) could have replaced the first one (extinct population) only very recently. Moreover, some cave populations could have evolved from the first extinct surface population recently too, but of course before its extinction. Very recently too, other cave populations could have evolved from the extant surface population. In summary, under the hypothesis of replacement of a surface population by another one, the coalescence time of mtDNA can give, at best, the older age possible for cavefish descendants of the extinct surface fish. Nevertheless, even if all cave settlements are very recent it does not preclude finding fish carrying divergent mtDNA.

Another study using a partial sequence of the cytochrome b gene confirmed the existence of two divergent mtDNA haplogroups [36]. This result was expected as mitochondrial genes are completely linked and a unique phylogeny is expected for mitochondrial genomes. Moreover, in this study a third haplogroup was identified in Yucatan. Using a more comprehensive sample and the same mtDNA marker [37], up to seven divergent haplogroups were found in Mexico (A to G, haplogroup G for cytb corresponding to haplogroup B with ND2) with a highly structured geographic distribution suggesting past fragmentation and/or a strong isolation by distance. In this study, haplogroup G was still cave specific (Piedras, Sabinos, Tinaja and Curva that are caves close to each other) and haplogroup A was still Northern Gulf coast and cave specific (**Figure 1**). However a more recent analysis [42], expanding further the sample of populations, allowed the identification of surface fish with haplotypes very close to haplogroup G (named Clade II lineage Ie) and haplogroup A (named Clade I Ia) in sympatry in a same water bodies, *i.e.* Mezquital and Aganaval, in Northwestern Mexico. This finding invalidates the hypothesis that haplogroup G evolved in the El Abra region a long time ago, went extinct, and was replaced by haplogroup A. Moreover, haplotypes “G like” were also found in surface fish localities (Rascon and Tamasopo) close to Sierra de El Abra [42].

Haplogroups A and G are highly divergent, supporting a model in which they accumulated mutations in two populations isolated for a long period of time. The presence of both haplogroups in Northwestern and Northeastern Mexico suggest that these populations mixed recently, at the time of a secondary contact. Dating results discussed below suggest that during the last glaciation, two allopatric populations from north Mexico, one carrying haplotypes belonging to haplogroup A and the other carrying haplotype belonging to haplogroup G, might have moved south and mixed there. After this glaciation they might have moved north again, now sharing haplotypes belonging to haplogroup A and G (this haplotype mixture is actually observed in the northwestern region, *i.e.* Mezquital and Aganaval water bodies). In the northeastern region, haplotypes belonging to the haplogroup G have up to now been found only in several caves in a restricted geographic area and haplotypes “G like” in surface localities also in a restricted area. Noteworthy, such recent secondary contact of divergent haplogroups were also observed at several other places in south Mexico [34, 42] suggesting that several populations of *Astyanax mexicanus* were isolated for a long time in different regions in Mexico and Central America and have been recently undergoing secondary contact.

In summary we think that, considering the mtDNA polymorphism alone, there is no reason to believe that the coalescence time of the mtDNA haplogroups should correspond to the age of the most ancient cavefish populations. On the contrary, taking into account the most recent publications, it suggests a recent admixture of two divergent populations. This admixture should be recent enough to allow the maintenance of both haplogroups at different geographic scale (in north Mexico as a whole and in northwestern and northeastern Mexico independently), as genetic drift should have eliminated, at a small geographic scale, one of them after a long period of time.

If we expand the phylogenetic analysis to nuclear sequences (nuDNA), the existence of two fish lineages implies that the mtDNA phylogeny should be congruent with unlinked nuDNA phylogenies, whereas a recent admixture of two surface populations before fish settlements in caves should lead to random fixation of alleles at different unlinked loci and thus incongruent phylogenies between mtDNA and nuDNA loci as well as between different unlinked nuDNA loci.

We compared mtDNA and nuDNA phylogenies using published sequences of several nuclear genes. First, we reconstructed a maximum likelihood phylogeny with Rag1. The resolution is so low that it precludes any phylogenetic inference, and even species defined using mtDNA are not supported. The congruence of the phylogenies with mtDNA and mtDNA+Rag1 [42] is the result of the very low quantity of phylogenetic signal in Rag1 compared to mtDNA and it does not support the congruence of mtDNA and Rag1 phylogenies.

Then we examined phylogenies obtained with other nuclear genes (Mc3r, Mc4r, Lepb, Lepr, Pomcb). These phylogenies are based on four sequences, but there are nevertheless highly informative. For each gene we found a unique phylogeny without homoplasy suggesting no recombination within each locus. Moreover, the incongruence of the phylogenies obtained with these unlinked loci supports their independent evolutionary histories. These results do not support two well defined fish lineages whereas admixture of gene phylogenies is expected when sampled localities are poorly isolated and/or have been separated for a short period of time.

Partial or complete coding sequences of five other genes (Per1, Per2, Tef1, Cry1a and Cpd photolyase) from three localities (Pachón and Chica caves and a surface locality close to Micos) were also available [52]. These sequences were aligned with the sequence of the Texas surface fish and the most parsimonious tree reconstructed for each gene. Surface fish sequences were always very close confirming the mtDNA evidence that the surface population sampled in Texas is genetically very close to Sierra de El Abra surface fish. These phylogenies are also interesting because they highlight another fact. Whereas for some genes, all the haplotypes are almost identical (very few mutations in Mc1r, Mc4r, Lepb, Pomcb, Per2, Tef1, Cpd photolyase), we can identify two, and only two, divergent haplotypes for Mc3r, Lepr, Per1 and Cry1a. Moreover, the distribution of divergent haplotypes is not the same for different loci (a divergent haplotype of Mc3r is found in Tinaja cave only, a divergent haplotype of Lepr in Molino cave only, divergent haplotypes of both Per1 and Cry1a at the surface only). Taking into account the existence of two divergent mtDNA (“G” haplotype in Tinaja and “A” haplotype in Pachón and surface fish), these phylogenies suggest that two divergent fish lineages with well differentiated genomes mixed and divergent alleles at each locus segregated randomly at the surface and in caves. When there are no divergent alleles at a locus (Mc1r, Mc4r, Lepb, Pomcb, Per2, Tef1, Cpd photolyase), one can suppose that alleles from one ancestral population went extinct. On this basis we came to the conclusion that cavefish could be much more recent than usually thought. In order to make a quantitative analysis of this hypothesis we applied two different approaches to estimate the age of some cave populations using multiple unlinked nuclear loci.

### Dating isolation times of cave populations

Dating the age of a recently isolated population that can exchange migrants with the “source” population is a difficult task [57]. If divergence is low and there is shared polymorphism between two populations, it can be the result of regular migration between these populations that diverged a long time ago, or the consequence of a recent divergence of completely isolated populations, or something in between. One can thus estimate how long ago the populations diverged (assuming no gene flow) using phylogenetic methods, or one can estimate the gene flow (assuming that the populations are at mutation/migration/drift equilibrium, *i.e.* they have been separated for a very long time, migrations occurred regularly and thus the phylogenetic signal has been erased) using population genetic methods. Such methods have been applied to study the evolution of *A. mexicanus* cavefish [32–39, 42, 43, 58], but neither one is of much use and often misleading if the goal is to develop a full picture that includes estimates of recent separation time and gene flow [57]. In such case it is necessary to consider non-equilibrium models and methods allowing the joint estimation of demographic parameters (populations sizes and migration rates) and divergence time [57]. Accordingly, we used IMa2, a widely used program for “isolation with migration” (IM) model analyses [53] to estimate divergence time between surface and cave populations with a dataset of multi-locus microsatellite polymorphism. IMa2 is based on backward simulations of coalescence of samples of alleles. For Pachón cave population, we estimated the divergence time using an alternative approach based on forward simulations of evolution of SNPs. Analyses of microsatellite polymorphism with IMa2 supported a recent origin of all cave populations. Analyses of SNPs confirmed a recent origin of Pachón cavefish.

### Dating with microsatellites

The microsatellite data set was kindly provided by M. Bradic and R. Borowsky [33]. We performed a series of pairwise analyses implying a cave population and a surface population or two cave populations using IMa2 in order to estimate the marginal posterior probability density of model parameters (*i.e.* population sizes, migration rates and divergence time). Whereas the current version of IMa2 can handle more than two populations, the phylogenetic relationships between populations must be known. In the case of *Astyanax mexicanus*, as shown above, there is no obvious phylogenetic relationships between surface and cave populations. In addition, as the number of parameters increases very fast with the number of populations analyzed, it is a very difficult task to analyze more than two populations. However, it is possible to study a large complex divergence problem that involves multiple closely related populations by analyzing pairs of populations [53, 59].

First we focused on the divergence time of Pachón cave population (O1, according to Bradic et al. nomenclature [33]) and a sample of surface fish from close localities (S3) (**Figure 1**) because large samples of alleles had been genotyped for many loci and it may allow more accurate estimations of model parameters than for cave populations for which a limited number of alleles were genotyped. Assuming a mutation rate, estimations of the effective population sizes, migration rates and divergence time can be obtained. In most population genetic studies, including *A. mexicanus* [33], the mutation rate of microsatellite loci is assumed to be about 5 × 10^−4^. This is a quite high mutation rate, but it is very likely as the loci retained for population genetic analyses are the most variable, thus those with the highest mutation rate. The estimation of the effective population size (N_e_) was 150 [150 – 1150] and 10,750 [6,650 – 16,350] for Pachón cave population and surface fish population, respectively. These estimations make sense as it is obvious after several trips in Sierra de El Abra that the census population size (N_c_) of surface fish is much higher than the census population size of the cavefish. N_c_ has been estimated for Pachón cave (8502; 95% confidence limits [1,279 – 18,283]) [1] but it has never been estimated for surface fish. On the one hand we expect that N_e_ is correlated with N_c_, but for fish that can potentially lay or fertilize hundreds of eggs or no eggs at all during their life such as *A. mexicanus* fish, the variance of the number of descendants is probably high (in the range of 10^1^ to 10^2^) and thus N_e_ might be one to two orders of magnitude smaller than N_c_ [60, 61]. If N_c_ is in the order of magnitude of 10^4^, it is expected that N_e_ is in the order of magnitude of 10^2^ to 10^3^ as found in the present study with IMa2. Previous studies, using Migrate [62], found similar N_e_ for several caves, including Pachón [33, 35].

The results obtained with IMa2 suggested that the migration rate from the cave to surface is negligible whereas the migration rate from the surface to the cave is low. This is also expected. If fish could easily exit the cave, no evolution of cavefish would have occurred. Moreover, it is difficult to imagine that blind cavefish who found their way to the surface will have a good fitness there. Concerning the migration rate of surface fish into the cave, it is likely that the fitness of surface fish in a cave is low compared to well-adapted cavefish (Luis Espinasa, personal communication). So even if the migration rate of adult surface fish into cave is not negligible, the “effective migration rate”, that is the rate of migration of surface fish that actually reproduce in the cave is probably extremely low. Accordingly, surface fish has never been observed in Pachón cave, but fish that were likely hybrids were reported during a couple of years in the 80’s [63]. This observation made by only one group of investigators had never been made by anybody else, before or after.

The IMa2 estimation of divergence time of 5,110 years [1,302 – 18,214] suggests that the Pachón population is much younger than usually thought. Indeed, and without any complex computations, a simple glance at the microsatellite data set actually supports a recent origin of this cavefish: 1) there is no divergence of the distribution of allele sizes found in the surface fish and Pachón cave fish, 2) the alleles found in the cave are also present at the surface. The difference between these populations is that, at each locus, there are many different allele sizes at the surface but a much smaller number of allele sizes in the cave. This differentiation without divergence can be easily explained by a much higher genetic drift in the small cave population than in the large surface population. Of note, even if we consider that the mutation rate is 10 times lower than the rate taken to make these estimations, the origin of the Pachón cavefish would still be much more recent than usually thought.

Similar results were obtained with all the cave populations studied. They all appeared having low N_e_ and a recent origin, less than 10,000 years ago. Taking into account the large variance of the divergence time estimations, we can conclude that they are most likely all less than 20,000 years old. In addition, taking also into account the uncertainty on the mean mutation rate of the microsatellites that could be about 5 times lower than assumed, the limit to the estimation of the age of the cave populations could be pushed to about 100,000 years.

### Dating with SNPs

Even though the recent evolution of *A. mexicanus* cavefish is well supported by analyses of multiple microsatellite loci with IMa2, this estimation was so at odds with the current opinion of antiquity of most cave populations that we considered necessary to bring additional evidence using a completely different approach and a totally different set of data. A congruent estimation of the model parameters of interest, in particular the divergence time, would greatly strengthen our conclusion. We focused on dating Pachón cave population for which we identified a large sample of SNPs in RNA-seq of pooled embryos of fish maintained in the lab. We performed the dating of the divergence time of this population with a Texas surface population of fish for which SNPs were also identified using the same approach. As discussed above, all data (mtDNA and nuDNA) showed unambiguously that these surface fish are closely related to the surface fish sampled in the Sierra de El Abra region. Despite this evidence and the absence of a high structuration of the genetic diversity of the surface fish in the Sierra de El Abra region, if someone is nevertheless convinced that there is a genetically closer surface population living near the Pachón cave, the straightforward conclusion is that the result we obtained is an overestimate of the age of the Pachón cave population. Of note, the surface fish shared about 7% of their SNPs with Pachón cavefish. As shared polymorphism is expected to decrease quickly if at least one population has a low effective population size and the two populations are completely isolated, it suggests that the divergence is recent or the migration rate is high.

In order to make estimations of the model parameters (*i.e.* population sizes, migration rates and divergence time) that could explain the distribution of SNPs within and between populations, we ran simulations of the evolution of the SNPs forward in time in two populations, allowing migration from the surface to the cave. Running these simulations with different sets of parameters allowed finding simulations for which, after a number of years of divergence, the distribution of the SNPs within and among simulated populations was very similar to the distribution observed within and among the real populations. The time of divergence was taken as an estimation of the age of the cave population. The use of simulations in order to estimate unknown values of model parameters is common in population genetics, for example in Approximate Bayesian Computation methods [64]. The rationale of this analysis is that when a population splits into two, genetic variation continues to be shared by the daughter populations for a period of time thereafter, even in the absence of gene exchange. As divergence proceeds, loci that were polymorphic in the ancestral population experience fixation of alleles in the descendant populations, and this sorting of alleles is part of the way the populations become different. Moreover, new alleles appear at new polymorphic sites and migration allows sharing of these new alleles too. As already discussed it is thus challenging to estimate if shared polymorphism is due to a recent split, gene flow or both [65]. However, some observations on SNPs distribution within and among the cave and surface populations suggested a recent divergence.

We found 2.34 times more derived alleles that reached fixation (substitutions) at synonymous polymorphic sites in the cavefish population than in the surface fish population. An excess of non-synonymous and non-coding substitutions were also observed. We searched for an explanation for these observations that could appear at first glance unexpected, in particular for synonymous substitutions which are for most of them probably neutral or nearly neutral. It is well known that if two populations have diverged for a long time and if the mutation rate is the same in both populations, the neutral substitution rates should be equal and independent of the population sizes [66]. Nevertheless, a simple explanation, totally compliant with this fundamental result of theoretical population genetics, relies on the fact that when an ancestral population is divided into a large (surface) and a small (cave) population, the probability of fixation of a neutral allele is the same in both populations (it is the allele frequency) but the process of fixation is faster in the small than in the large population. Indeed the mean time to fixation 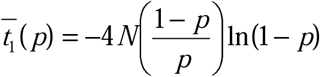 (where *N* is the population size, *p* is the allele frequency) [67]. The straightforward consequence is a transient acceleration of the substitution pace in the small population that is not anymore observed after a long period of time [66]. We thought that this information, together with other information about the distribution of polymorphism within and between populations, could be used for divergence dating.

We defined summary statistics describing the polymorphism and the divergence of two populations that could be accurately estimated using pooled RNA-seq [68]. The evolutionary model is identical to the model analyzed with IMa2 and relies on the same parameters (three effective population sizes, two migration rates, and a divergence time). Of note, we set the migration rate from cave to surface to zero as the analyses with IMa2 and several trips to this cave convinced us that the impact of migration of Pachón cavefish on surface fish DNA polymorphism is negligible.

In our simulations, we set the generation time to two and five years for the surface and cave populations, respectively. This surface fish generation time is twice the estimations obtained for other *Astyanax* species [69] and the cavefish generation time is the value estimated by P. Sadoglu, unpublished but reported as a personal communication [40]. This estimation is based on the hypothesis that cavefish may live and remain fertile for a long time, about 15 years. It is unlikely that these generation times are underestimates and they could actually be overestimates of the true generation times. As the estimation of the age of the Pachón cavefish population directly depends on these generation times, the divergence times we discuss below are more likely overestimates than underestimates. We took into account migration rate from surface to cave. This migration rate depended on two parameters: the probability of migration/year and the percentage of cavefish that were surface migrants when a migration occurred. This way it is possible to simulate different possibilities such as few migrants that enter the cave very often or many migrants in very rare occasions, and all intermediate cases between these two extremes.

We also took into account that genetic drift occurred during several generations in the lab. In order to estimate the effect of genetic drift in the lab and the accuracy of alleles frequency estimations using pooled RNA-seq, we compared the frequencies of two alleles (Pro106Leu) of MAO gene estimated with a sample of wild caught Pachón fish, a sample of adult fish maintained in the lab and estimated using pooled RNA-seq of lab embryos. Although it has been published that Pachón population is not polymorphic at this locus [70], we found polymorphism with a larger sample of wild caught fish and similar frequencies with both adult and embryo lab fish (data not shown) suggesting that genetic drift did not remove from the lab population a polymorphism known to be present in the Pachón population.

Of note whereas the estimation of the divergence time depends on the generation time and the mutation rate as with IMa2, our estimation depends only on the generation time. This is due to the fact that the summary statistics we used, *i.e.* SNPs class frequencies, to compare simulated and real populations, do not depend on the mutation rate. We set a mutation rate allowing a number of SNPs within and between simulated populations large enough to have accurate estimations of the summary statistics.

Without migration, shared polymorphisms were quickly lost and the best fit of the model to the data was obtained when the cavefish population size (1,250) was smaller than the surface fish population (10,000) and the age of the cavefish population was 20,600 years. When migration was included, good fit with the data also implied large differences in population sizes, a low migration rate and low numbers of migrants. The very best fit was observed for a surface fish population size of 20,000, a cave population size of 1,250, and a cave population age of 54,200 years. Nevertheless, most very good fits were obtained with a divergence time in the range of 20,000 to 30,000 years.

As expected no fit was found when the surface and cave populations had similar sizes. The best fit was observed when the ancestral population size was similar to the surface population size. It is at odds with the estimation of a much larger ancestral population size estimated with IMa2. We do not have a clear explanation for this discrepancy, but it could be the consequence of the admixture of two divergent populations at the origin of the ancestral population. Such admixture could have increased the number of alleles at each microsatellite loci, and IMa2 thus inferred a larger ancestral population size. For SNPs, admixture of divergent populations results in more polymorphic sites, but the number of alleles at a given locus is not increased as the probability that two different mutations occurred independently at the same locus in two populations is extremely low.

### Other evidence for a recent origin of the Pachón cavefish

In a recent analysis of the expression of 14 crystallin genes in the Pachón cavefish, 4 genes are not expressed or expressed at a very low level, but no stop codon or frameshift could be identified [26]. This result is in accordance with a recent origin of this population, as several loss-of-function mutations should have reached fixation after several hundred thousand years of evolution of genes that would no longer be under selection, as they are not necessary in the dark [47]. Indeed, other fish species that are likely confined into caves for millions of years have fixed loss-of-function mutations in several opsins and crystallins genes [48–50]. We are currently working on dating cavefish populations using the frequency of loss of function mutations in genes that are dispensable in the dark.

Second, a recent study has shown that the heat shock protein 90 (HSP90) phenotypically masks standing eye-size variation in surface populations [71]. This variation is exposed by HSP90 inhibition and can be selected for, ultimately yielding a reduced-eye phenotype even in the presence of full HSP90 activity. This result suggests that standing genetic variation in extant surface populations could have played a role in the evolution of eye loss in cavefish. This is also compatible with a recent origin of the cave population.

### Non-equilibrium models and cavefish population genetics

A recent origin of the so-called “old” Pachón population can solve a conundrum put forward by previous and the present analyses. First, at the SNP and microsatellite level, the diversity is not that low in Pachón cave when compared with surface populations, *i.e.* about one third. If the populations are at migration/drift equilibrium, it means that the effective population size of Pachón cavefish is about one third of the surface populations, and this is at odds with the likely large difference in census population sizes [1, 33]. Of course, we can propose *ad hoc* hypotheses to explain this discrepancy. Cavefish may have a much lower reproductive success variance than surface fish, or surface fish could have larger population size fluctuations through time than cavefish. In such cases, the effective population sizes could be much closer to one another than census population sizes because it is well established that large variance in reproductive success and large population size fluctuations hugely reduce the effective population size [61]. An alternative explanation is that the genetic diversity in the Pachón cave is actually higher than expected at mutation/drift/migration equilibrium. Our results suggest that the effective population size of the surface fish is at least one order of magnitude larger than the effective population size of cavefish, a ratio that is more in accordance with the unknown but certainly very different long term census population sizes. The present study is a striking illustration of how misleading analyses of evolutionary processes that do not take account that biological systems are not necessarily at equilibrium can be. The two analyses we described above rely on approaches that do not suppose mutation/migration/drift equilibrium. They allowed the estimation of demographic parameters that are more in line with expectations based on field observations than previous estimations. There are much more surface fish than cavefish and the impact of migration of cavefish on surface fish diversity is likely extremely low, and most likely null.

The new time frame we propose for the evolution of the Pachón cave population would not allow enough time for the fixation of many *de novo* mutations and most derived alleles that reached fixation in caves were probably already present in the ancestral population. This was also suggested by a recent population genomic study [72]. This may imply that the cave phenotype evolved mainly by changes in the frequencies of alleles that were rare in the ancestral surface population. In particular, some of these alleles would have been loss-of-function or deleterious mutations that could not reach high frequency in surface populations but they could reach high frequency or fixation quickly in a small cave population where they are neutral or even advantageous.

Although we estimated that all cavefish populations are probably recent, less than 20,000 years old, the number of independent and approximatively simultaneous adaptations to cave and evolution of cave phenotype is still an open question. The evolution in a short period of time of the phenotype of individuals belonging to a population adapting to a new environment, is actually not that unexpected and has already been observed in other fish species such as the stickleback [73], dwarf whitefishes [74] and African cichlids [75, 76].

Recently, the first European cavefish, with a well differentiated cave phenotype, has been described. The phylogenetic analysis of mtDNA haplotypes, the analysis of genetic differentiation using microsatellite loci and the recent glacial history of the region suggests that these fish population is highly isolated but for less than 20,000 years [77].

Mexican cavefish could thus be another and striking illustration that many phenotypic changes can accumulate in parallel and in a short period of time thanks to standing genetic variation [78]. The relative roles of selection and drift in allelic frequency changes is not yet understood, but if the recent origin of cavefish populations is confirmed, they would be an excellent model to analyze this issue using population genomics tools such as the quantification of selective sweep around candidate loci most likely involved in the adaptation to a cave environment.

## Materials and Methods

### Dating with IMa2

With a multilocus microsatellite polymorphism dataset [33] and the program IMa2 [53] divergence times of pairs of populations were estimated. The program is based on an isolation-with-migration (IM) model and uses Metropolis-coupled Markov chain (MCMC) techniques to estimate the posterior densities of the time of divergence, population sizes and gene flow [57]. The model assumes random population samples, a stepwise mutation model, neutral mutation, freely recombining loci and constant population sizes and gene exchange rates. Although modelling constant population sizes and gene exchange rates might not be ideal in the case of cavefish populations, modelling general patterns requires simplifications and this is presently the only option in programs such as IMa2 and Migrate which is the program previously used to estimate gene flow [33, 62]. It was also not possible to take into account changes in generation time after the separation of the populations. It is thus more the order of magnitude of the parameter estimations that can be discussed rather than maximum likelihood values *per se*. As the upper limit of the prior distribution of each model parameter must be set, we ran short MCMC chains to test several sets of parameters that allowed the identification of suitable values of upper limits mutation-scaled effective population sizes, gene flows and divergence time. It allowed also to estimate the length of the burn-in period. In order to use a large and even number of alleles/locus/populations we sampled at random the same number of alleles at each locus in each population. Moreover, we restrained the number of allele to 60 as it took more than one month to get the results with 22 loci and 60 alleles/locus/population. Such a large sample of alleles for many loci was not available for many comparisons. In that cases, we used smaller samples of loci and allele/locus. Nevertheless, we did retain analyses with less than 20 alleles/locus as the results were not robust according to a resampling process. The parameter sets used in the different analyses are indicated as a command line in the legend of the figures that summarize the output of IMa2. The number of loci and the number of alleles/locus/population are also indicated (**Additional file 1; Figure S3 to Figure S9**).

### Sampled populations for SNPs analyses

For fifteen years we have maintained laboratory stocks of *Astyanax mexicanus* cavefish and surface fish, founded with fish collected respectively in the Pachón cave (Sierra de El Abra, Mexico) and at the San Solomon Spring (Texas, USA) (**Figure 1**), and obtained from W. R Jeffery in 2004. In 2012, we purchased thirty *Hyphessobrycon anisitsi* fish.

### RNA samples and RNA-seq

In order to identify polymorphisms at the population level based on a Pool-seq approach [68], for each population, 50 to 200 embryos/larvae from several independent spawning events and at different developmental stages (6 hours post-fertilization to two weeks post-fertilization) were pooled and total RNA isolated. (**Additional file 1; Figure S10**). Each RNA sample was sequenced on an Illumina HiSeq 2000 platform (2 × 100 bp paired-end). The pooled embryo samples had been previously sequenced using the Sanger and 454 methods [79] (**Additional file 1; Figure S10**).

### Transcriptome assembly and annotation

The *Astyanax mexicanus* transcriptome was assembled with Newbler ver. 2.8 (Roche 454) sequence analysis software using 454 sequences (2.10^6^ reads) of both the Pachón cave and surface fish pooled embryos. We obtained 33,400 contigs (mean contig length = 824 bp). We also tried to generate a transcriptome assembly using the Illumina sequences, but whereas this resulted in more contigs (49,728) than the 454 sequences, many of them were concatenations of different transcripts and in some cases the same transcript was found in more than one contig. We therefore mapped the Illumina sequences onto the 454 contigs to identify and annotate SNPs. Putative coding sequences in each contig were identified using the zebrafish (Zv9) proteome available at EnsEMBL 73 as a reference [80]. A contig was considered protein coding if the e-value for the best hit was < 10^−5^. We found 13,240 protein coding contigs (contig mean length = 530 bp). We identified contigs containing domains that matched different zebrafish proteins and which were most likely chimeric contigs. These contigs were removed (369, *i.e.* 3% of the protein coding contigs). In total, we analyzed 12,871 putative protein coding contigs.

### SNP identification and annotation

Illumina sequences were aligned to contigs with BWA [81] using the default parameters for paired-end reads. *Hyphessobrycon anisitsi* sequences were aligned to *Astyanax* contigs using a lower maximum edit distance (n = 0.001).

SNPs calling was performed using GATK UnifiedGenotyper v2.4.9 [82]. Because we filtered SNPs after detection using different parameter thresholds described below, we used the allowPotentiallyMisencodedQuals and –rf BadCigar options. We detected 299,101 SNPs including 141,490 SNPs in annotated contigs.

When a complete coding sequence was identified, *i.e.* from the start codon to the stop codon and corresponding to a complete zebrafish protein, we could identify the non-coding flanking sequences (containing 18,743 SNPs), otherwise only the sequence matching the coding sequence of the zebrafish was annotated as coding and the flanking sequences were not annotated. The 55,950 SNPs in the coding sequences were annotated as synonymous or non-synonymous, according to which amino acid was coded for by the alternative codons resulting from the SNP. The ancestral allele and the derived allele were inferred according to the allele found in the outgroup *Hyphessobrycon anisitsi* (**Figure 4**). SNPs for which the ancestral allele and derived allele could not be identified, either because in *Hyphessobrycon anisitsi* no sequence could be identified or there was another allele present or the allele was polymorphic, were discarded.

### SNP classification

The SNPs identified in *Astyanax mexicanus* SF and CF were classified into eight classes (**Figure 4**). The number of SNPs in the different classes depended on the thresholds used to consider a SNP as reliable and polymorphic in each population. The rationale for the set of thresholds selected is given below.

The populations being closely related (they belong to the same species) and the mutation rate for a SNP origin being very low (~10^−8^), we would expect that the eighth class (divergent polymorphism) of SNPs would be a very rare outcome because it is the result of two independent mutations at the same site, either in the ancestral population or in the CF and SF populations. We found only one SNPs in this class **(Table 3)**. It suggests that Illumina sequencing did not generate a number of sequencing errors that would significantly inflate the number of SNPs identified.

### Parameter thresholds for SNP selection

We examined the effect of the thresholds applied to parameters used to discard SNPs before their classification and population genomics analyses.

First we looked at the effect of sequencing depth. Whereas the mean sequencing depth was 820, the standard deviation was very large (9,730). When the minimal number of reads per population at a SNP site was set to 100 or higher, the relative frequencies of the eight SNP classes were very stable, indicating that 100 was a good compromise between the stability of the distribution of the SNPs into different classes and the number of SNPs discarded (**Additional file 1; Figure S11**).

We then considered the effect of the e-value of the blast between the *Astyanax* contig and the zebrafish sequence used for annotation, in order to discard poorly conserved sequences that were misidentified as protein coding. It appeared that the SNP classification was stable whichever the threshold was used, *i.e.* e-value < 10^−5^ (**Additional file 1; Figure S11**).

We also examined the effect of the interval between SNPs, because we would expect clusters of spurious SNPs in poorly sequenced regions. We tested the effect of selecting SNPs in regions without any other SNPs. As expected, there was an excess of shared polymorphisms (class 7) with a small window size. When the threshold was set to > 50 bp on each side of the SNP, the distribution was stable (**Additional file 1; Figure S11**).

Finally, we considered that the lowest value of minor allele frequency (MAF) in the lab populations should be set around 5% because the effective population size in the lab is low. All the above thresholds, apart from that for MAF, are trade-offs between quality and quantity of the data. The lowest MAF value possible in the pooled embryo samples depends on the unknown number of parents of the embryos, and the MAF threshold of >5% could therefore be considered arbitrary. Nevertheless, using MAF thresholds of 1%, 5% and 10% we obtained similar SNP class frequencies (**Table 3** and **Additional file 1; Table S2a** and **Table S2b**).

The results were thus also robust according to this parameter, and the use of different sets of parameters led to similar distribution of SNP classes that led to the same conclusion.

Therefore, all analyses in this paper were performed using the following thresholds: MAF > 5%; depth > 100; e-value < 10^−5^; SNP isolation > 50 bp.

### Simulations of the evolution of neutral polymorphisms in two populations

In order to estimate the age of the Pachón cave population, we compared the distribution of SNPs into seven classes (the divergent polymorphism class was empty and thus excluded) defined above with the distribution obtained in simulations of the evolutionary process (**Figure 3**). More detailed on the rationale of the method is given in **Additional file 4**. The evolutionary model and its parameters are those of IMa2, *i.e.* effective population sizes, migration rates and divergence time. The range of values tested were defined according observation in the field, estimations found in the literature and the results of the analysis with IMa2. The full model is as following: an ancestral population with a given size (10,000, 20,000, 50,000 or 100,000) and at mutation/drift equilibrium (which depends on the mutation rate and the population size) was split into two populations that could have different sizes. There could be migrations from the surface to the cave. The smallest probability of migration was set to 0.00001 and the highest to 0.1 per year. The smallest percentage of cavefish that are migrants when a migration occurs was set to 0 and the highest was set to 1%. We also took into account that genetic drift could have occurred in the laboratory stocks (effective population size in the lab was set to 10 and the number of generation was set to 10). All mutations were neutral (frequency changes at each locus were driven by genetic drift only) and each locus were evolving independently. For a given set of parameters, each ten generations after the isolation of two populations, we estimated the frequency of SNPs in each category and we estimated a score of goodness of fit with the frequencies obtained with the real SNP data set. We ran the simulation and the test of goodness of fit with different sets of parameters in order to identify the sets of parameters, including the age of the Pachón cave population, that resulted in SNP class frequencies that fitted well with observed frequencies. The program was written in C and is available on Github with its documentation (http://julienfumey.github.io/popsim).

### Data storage and analyses

SNPs and their annotations are stored in a MySQL database and are available online at http://ngspipelines.toulouse.inra.fr:9022. Perl and R scripts for the data analyses and graphics are available upon request. Illumina and 454 transcriptomic sequences are available on ArrayExpress: https://www.ebi.ac.uk/arrayexpress/experiments/E-MTAB-5142/

## Additional Material

**Additional file 1:** Supplementary figures and tables

**Additional file 2:** Four taxon unrooted nuclear gene phylogenies

**Additional file 3:** Allele frequency distributions at 25 microsatellite loci

**Additional file 4:** Rationale of dating with pool-seq SNPs

## Acknowledgments

This work has benefited from the facilities and expertise of the high throughput sequencing platform of I2BC. The work was supported by a collaborative ANR (Agence Nationale de la Recherche) grant BLINDTEST and IDEEV.

### Authors’ contributions

DC and SR designed the study. JF wrote the program of simulation. DC and JF analyzed the data. DC, CT, SR, HH collected the data. CN and JF generated the databases. DC drafted the manuscript. All authors contributed to the writing of the manuscript. All authors read and approved the final manuscript.

### Competing interests

The authors declare that they have no competing interest.

### Animal ethics

SR’s authorization for use of animals in research is number 91-116, and includes a “Certificat de capacité pour l’élevage de faune sauvage”. Experiments were performed according to Paris Centre-Sud Ethic Committee authorization numbers 2012-0053 and 2012-0056. Fin-clips from wild-caught animals were collected under the auspices of Mexican permit 02241/13 delivered to SR by Secretaria de Medio Ambiente y Recursos Naturales.

